# SARS-CoV-2, an evolutionary perspective of interaction with human ACE2 reveals undiscovered amino acids necessary for complex stability

**DOI:** 10.1101/2020.03.21.001933

**Authors:** Vinicio Armijos-Jaramillo, Justin Yeager, Claire Muslin, Yunierkis Perez-Castillo

## Abstract

The emergence of SARS-CoV-2 has resulted in more than 200,000 infections and nearly 9,000 deaths globally so far. This novel virus is thought to have originated from an animal reservoir, and acquired the ability to infect human cells using the SARS-CoV cell receptor hACE2. In the wake of a global pandemic it is essential to improve our understanding of the evolutionary dynamics surrounding the origin and spread of a novel infectious disease. One way theory predicts selection pressures should shape viral evolution is to enhance binding with host cells. We first assessed evolutionary dynamics in select betacoronavirus spike protein genes to predict where these genomic regions are under directional or purifying selection between divergent viral lineages at various scales of relatedness. With this analysis, we determine a region inside the receptor-binding domain with putative sites under positive selection interspersed among highly conserved sites, which are implicated in structural stability of the viral spike protein and its union with human receptor hACE2. Next, to gain further insights into factors associated with coronaviruses recognition of the human host receptor, we performed modeling studies of five different coronaviruses and their potential binding to hACE2. Modeling results indicate that interfering with the salt bridges at hot spot 353 could be an effective strategy for inhibiting binding, and hence for the prevention of coronavirus infections. We also propose that a glycine residue at the receptor binding domain of the spike glycoprotein can have a critical role in permitting bat variants of the coronaviruses to infect human cells.

## Introduction

The recent emergence of the novel SARS coronavirus 2 (SARS-CoV-2) marked the third introduction of a highly pathogenic coronavirus into the human population in the twenty-first century, following the severe acute respiratory syndrome coronavirus (SARS-CoV) and the Middle East respiratory syndrome coronavirus (MERS-CoV). The first, SARS-CoV emerged in November 2002 in the Guangdong province of China and spread globally during 2002-03, infecting more than 8000 people and causing 774 deaths (Drosten et al., 2003; WHO, 2004). MERS-CoV was the second emergence and was first detected in Saudi Arabia in 2012 and resulted in nearly 2500 human infections and 858 deaths in 27 countries (Fehr et al., 2017; Zaki et al., 2012). In December 2019, SARS-CoV-2, a previously unknown coronavirus capable of infecting humans was discovered in the Chinese city of Wuhan, in the Hubei province (Huang et al., 2020; Zhu et al., 2020). SARS-CoV-2 is associated with an ongoing pandemic of atypical pneumonia, now termed coronavirus disease 19 (Covid-2019) that has affected over 209,000 people with 8778 fatalities as of March 19, 2020 (WHO, 2020). Both SARS-CoV and MERS-CoV are thought to have originated in colonies of bats, eventually transmitted to humans, putatively facilitated by intermediate hosts such as palm civets and dromedary camels, respectively (Cui et al., 2019). The genome of SARS-CoV-2 shares about 80% nucleotide identity with that of SARS-CoV and is 96% identical to the bat coronavirus BatCoV RaTG13 genome, reinforcing the probable bat origin of the virus (Zhou et al., 2020). However, better assessing the evolutionary dynamics of SARS-CoV-2 is an active research priority worldwide.

SARS-CoV, MERS-CoV and SARS-CoV-2 belong to the genus *Betacoronavirus* within the subfamily *Coronavirinae* of the family *Coronaviridae*. Members of this family are enveloped viruses containing a single positive-strand RNA genome of 27-32 kb in length, the largest known RNA virus genome. The coronavirus spherical virion consists of four structural proteins: the spike glycoprotein (S-protein), the envelope protein, membrane protein and nucleocapsid. The transmembrane trimeric S-protein plays a critical role in virus entry into host cells (Gallagher & Buchmeier, 2001; Tortorici & Veesler, 2019). It comprises two functional subunits: S1 subunit, where the receptor-binding domain (RBD) is found, is responsible for binding host cell surface receptors and S2 subunit mediates subsequent fusion between the viral and cellular membranes (Kirchdoerfer et al., 2016; Yuan et al., 2017). Both SARS-CoV and SARS-CoV-2 interact directly with angiotensin-converting enzyme 2 (ACE2) to enter host target cells (Hoffmann et al., 2020; Li et al., 2003; Walls et al., 2020; Yan et al., 2020). In the case of SARS-CoV, ACE2 binding was found to be a critical determinant for the virus host range and key amino acid residues in the RBD were identified to be essential for ACE2-mediated SARS-CoV infection and adaptation to humans (Li et al., 2006; Li et al., 2005).

Understanding the dynamics that permits a virus to shift hosts is of considerable interest, and further be an essential preliminary step towards facilitating the development of vaccines and the discovery of specific drug therapies. We employ a multidisciplinary approach to look for evidence of diversifying selection on the S-protein gene, and model the interactions between human ACE2 (hACE2) and the RBD of selected coronavirus strains, which ultimately afforded us novel insights detailing virus and host cell interactions. Given the rapid pace of discovery we aim to add clarity to evolutionary dynamics of diseases strains by more precisely understand the dynamics at the S-protein and its interaction with hACE2.

## Methods

### Phylogenetic reconstruction, and analysis testing for evidence of positive/purifying selection in the coronavirus S-protein region

The most similar genomes to SARS-CoV-2 MN908947 were retrieved using BLASTp (Altschul et al., 1997) vs the NR database of GenBank (Table 1). Genomes were then aligned using MAUVE (Darling et al., 2004) and the S-protein gene was trimmed. The extracted genomic sections were aligned using a translation align option of Geneious (Kearse et al., 2012) with a MAFFT plugin (Katoh & Standley, 2013). The phylogenetic reconstruction of S-protein genes was performed with PhyML (Guindon et al., 2010), using a GTR+I+G model, using 100 non-parametric bootstrap replicates. Both, the alignment and the tree were used as input for PAML Codeml (Yang, 2007). The presence of sites under positive selection was tested by the comparison of M2 (it allows a proportion of positive, neutral and negative selection sites in the alignment) vs M1 (it allows a proportion of neutral and negative selection sites in the alignment) and M8 (ω follows a beta distribution plus a proportion of sites with ω>1) vs M7 (ω follows a beta distribution) models using the ETE toolkit 3.0 (Huerta-Cepas et al., 2010). The presence of tree nodes under positive selection was obtained with the free branch model and then tested by the comparison of branch free (different ω for each selected branches) vs M0 (negative selection for all sites and branches-null model) and branch free vs branch neutral (ω=1 for selected branches) models. The presence of sites with positive selection under specific branches of the tree was tested with bsA (proportion of sites with positive selection in a specific branch of the tree) vs bsA1 (proportion of sites with neutral and purifying selection in a specific branch of the tree) models. Likelihood ratio test (LRT) was performed (p≤ 0.05) to compare the hypothesis contrasted by each model.

**Table 1.** List of coronavirus isolates used for positive selection analysis (closer dataset)

| Accession number (NCBI) | Description |
| --- | --- |
| AY390556 | SARS coronavirus GZ02 |
| DQ071615 | Bat SARS-like coronavirus Rp3 |
| DQ084199 | Bat SARS-like coronavirus HKU3-2 |
| DQ412043 | Bat SARS-like coronavirus Rm1 |
| FJ211859 | Recombinant coronavirus clone Bat SARS-CoV |
| FJ211860 | Recombinant coronavirus clone Bat-SRBD |
| KC881006 | Bat SARS-like coronavirus Rs3367 |
| KF569996 | Rhinolophus affinis coronavirus isolate LYRa11 |
| KJ473814 | BtRs-BetaCoV/HuB2013 |
| KT444582 | SARS-like coronavirus WIV16 |
| KY417142 | Bat SARS-like coronavirus isolate As6526 |
| KY417144 | Bat SARS-like coronavirus isolate Rs4084 |
| KY417146 | Bat SARS-like coronavirus isolate Rs4231 |
| KY417148 | Bat SARS-like coronavirus isolate Rs4247 |
| MG772933 | Bat SARS-like coronavirus isolate bat-SL-CoVZC45 |
| MG772934 | Bat SARS-like coronavirus isolate bat-SL-CoVZXC21 |
| MK211375 | Coronavirus BtRs-BetaCoV/YN2018A |
| MK211378 | Coronavirus BtRs-BetaCoV/YN2018D |
| MN996532 | Bat coronavirus RaTG13 |
| MT084071 | Pangolin coronavirus isolate MP789 |
| MN908947 | Wuhan seafood market pneumonia virus isolate Wuhan-Hu-1 (SARS-CoV-2) |

We used the set of programs available in HyPhy (Kosakovsky Pond et al., 2020), Fast Unconstrained Bayesian AppRoximation (FUBAR) to detect overall sites under positive selection, and Fixed Effects Likelihood (FEL) to detect specific sites under positive selection in specific branches. We used Mixed Effects Model of Evolution (MEME) to detect episodic positive/diversifying selection and adaptive Branch Site REL (aBSREL) to detect branches in the tree under positive selection. The web server Datamonkey (Weaver et al., 2018) was used to perform the HyPhy analyses.

Finally, TreeSAAP 3.2 (Woolley et al., 2003) was used to detect sites under adaptation (in terms of physicochemical properties). The same alignment and tree described above were used for this analysis. All these experiments were performed again using the S-protein genes of a shorter list of accessions and more distantly related (broad dataset) to SARS-COV-2 (AY304488, AY395003, DQ412043, FJ882957, KY417144, MG772933, MG772934, MN908947, NC_004718) to test the reproducibility of the predicted branches and sites under positive selection.

### Assessing molecular structure and host-specific interactions

The crystal structure of the SARS-CoV S-protein RBD (GeneBank ID NC_004718) in complex with hACE2 was retrieved from the Protein Data Bank (code 2AJF) (Berman et al., 2000). Homology models were constructed using this structure as template for the RBDs of SARS-CoV-2 (SARS2, GeneBank ID MN908947), the Bat SARS-like coronavirus isolate Rm1 (Rm1, GeneBank ID DQ412043) and the Bat SARS-like coronavirus isolate Rs4231 (Rs4231, GeneBank ID KY417146). One additional homology model for the G496D mutant of the SARS-CoV-2 RBD (SARS2-MUT) was constructed. Homology models were built with Modeller v. 9 (Webb & Sali, 2016) using its UCSF Chimera interface (Pettersen et al., 2004). Five models were constructed for each target sequence and the one with the lowest DOPE score was selected for the final model.

All non-amino acidic residues were removed from the SARS-CoV RBD-hACE2 complex to obtain a clean complex. The homology models of the SARS2, Rm1, Rs4231 RBDs and SARS2-MUT were superimposed into the SARS-CoV RBD to obtain their initial complexes with hACE2. These complexes were then subject to molecular dynamics (MD) simulations and estimation of their free energies of binding using Amber 18 (Case et al., 2018). For the later, ACE2 was considered as the receptor and the RBDs as ligands. The protocol described below was employed for all complexes and otherwise noted default software parameters were employed.

Systems preparation was performed with the tleap program of the Amber 18 suite. Each complex was enclosed in a truncated octahedron box extending 10 Å from any atom. Next, the boxes were solvated with TIP3P water molecules and Na+ ions were added to neutralize the excess charge. Systems were minimized in two steps, the first of which consisted in 500 steps of the steepest descent algorithm followed by 500 cycles of conjugate gradient with protein atoms restrained using a force constant of 500 kcal/mol.Å^2^. The PME method with a cutoff of 12 Å was used to treat long range electrostatic interactions. During the second minimization step the PME cutoff was set to 10 Å and it proceeded for 1500 steps of the steepest descent algorithm followed by 1000 cycles of conjugate gradient with no restrains. The same PME cutoff of 10 Å was used in all simulation steps from here on. Both minimization stages were performed at constant volume.

The minimized systems were heated from 0 to 300 K at constant volume constraining all protein atoms with a force constant of 10 kcal/mol.Å^2^. The SHAKE algorithm was used to constrain all bonds involving hydrogens and their interactions were omitted from this step on. Heating took place for 10000 steps, with a time step of 2 fs and a Langevin thermostat with a collision frequency of 1.0 ps^−1^ was employed. All subsequent MD steps utilized the same thermostat settings. Afterward, the systems were equilibrated for 100 ps at a constant temperature of 300 K and a constant pressure of 1 bar. Pressure was controlled with isotropic position scaling with a relaxation time of 2 ps. The equilibrated systems were used as input for 10 ns length production MD simulations.

The free energies of binding were computed under the MM-PBSA approach implemented in AmberTools 18 (Case et al., 2018). A total of 100 MD snapshots were evenly selected, one every 50 ps, from the last 5 ns of the production run for MM-PBSA calculations. The ionic strength was set to 100 mM and the solute dielectric factor was set to 4 for all systems.

## Results and discussion

In order to detect branches and sites under positive/negative selection, two datasets were explored. The first (‘closer’ dataset) harbors the most similar genomes to Wuhan-Hu-1 coronavirus (SARS-CoV-2) (MN908947). For this dataset, several genomes were excluded from the analysis because they showed minimal variation to other sequences. We used a preliminary phylogeny to select a representative isolate of each clade (Table 1) in order to exclude highly similar sequences. The second dataset (‘broad’ dataset) includes some accessions of the first dataset plus isolates less related to SARS-CoV-2, like SARS-like coronavirus isolates from different countries (see methods). We compare the results of two dataset because the phylogenetic distance between orthologues in a given dataset has been demonstrated to alter the ability to detect selection in PAML and MEME (McBee et al., 2015).

In both datasets, we observed evidence of purifying selection in the majority of nodes of the tree. Specifically, in the ‘closer’ dataset we identified 38 nodes with evidence of negative selection, and 4 under positive selection when free ratios model of CODEML model was applied. To confirm the four nodes under positive selection we use LTR test for contrasting hypothesis using branch free, branch neutral and M0 models of CODEML. Using these approximations, any node predicted by free ratios model with ω>1 was significantly different to the purifying (ω<1) or neutral (ω=1) models. An equivalent analysis was performed using aBSREL of HyPhy, observing episodic diversifying selection in at least 8 of 41 nodes of the phylogenetic tree reconstructed with the ‘closer’ dataset (Figure 1). Interestingly, one of the divisions detected with diversifying selection was the branch that contains SARS-CoV-2, Pangolin coronavirus isolate MP789 and Bat coronavirus RaTG13 (called SARS-CoV-2 group) but not the specific branch that contains SARS-CoV-2.

**Figure 1.**
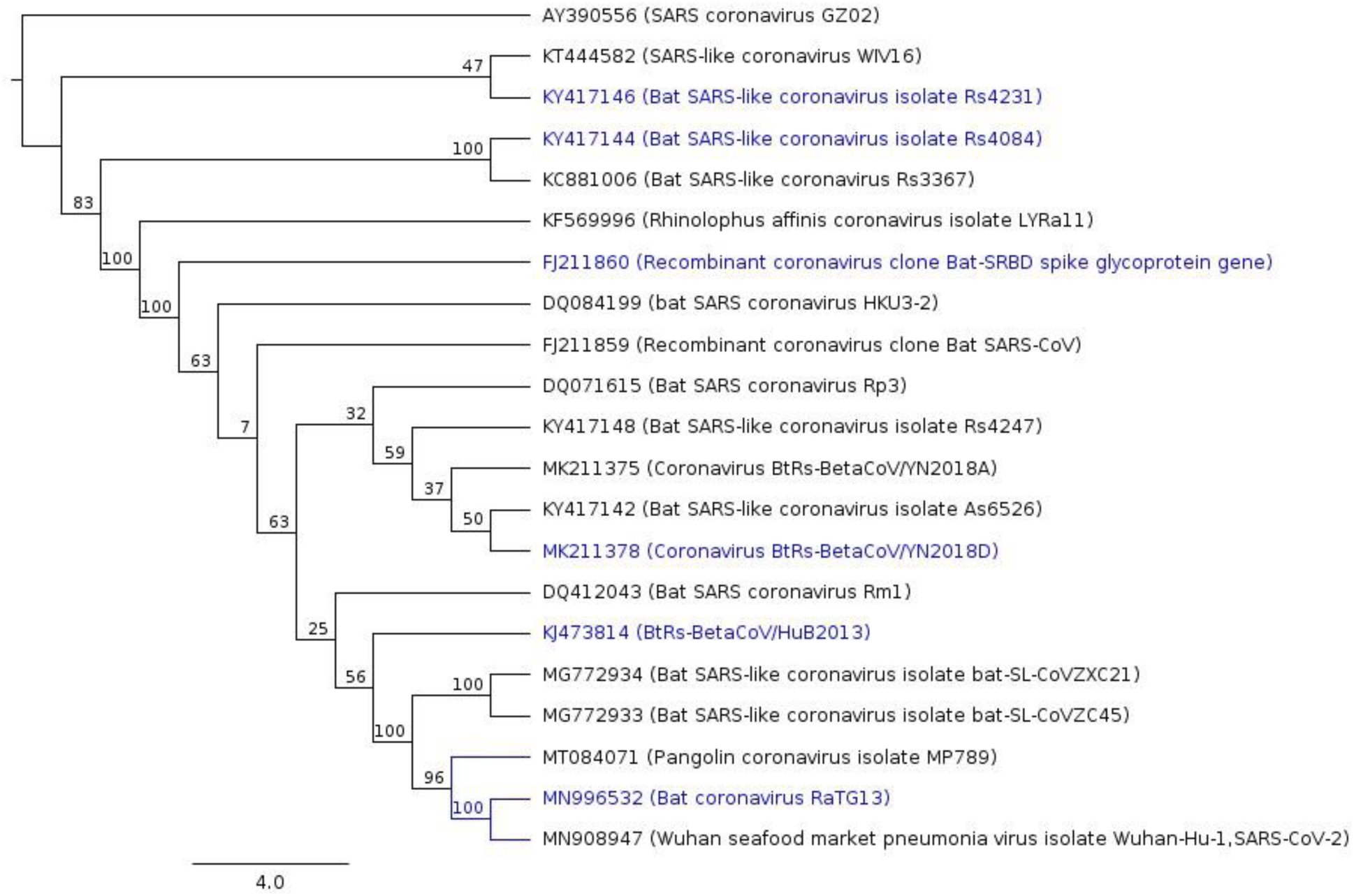
Phylogenetic tree of S-protein genes of selected betacoronaviruses (‘closer’ dataset). Predicted nodes under episodic diversifying selection (according to predictions generated by aBSREL of HyPhy) were colored in blue. Numbers in the inner nodes represent the number of non-parametric bootstrap replicates

MEME of HyPhy detected 40 sites in different branches with signal of episodic positive selection, but FEL did not identify sites under positive selection (with p < 0.1) in the SARS-CoV-2 when ‘closer’ dataset was used. Depending on the focal dataset analyzed (‘close’ vs. ‘broad’), FEL detected several sites under positive selection in SARS-CoV-2, demonstrating the influence of relatedness within the dataset in branch-sites model predictions. FEL predicted the site F486 under positive selection in SARS-CoV-2 using the closer dataset without Pangolin coronavirus isolate MP789. It is interesting despite the influence of the dataset in the results, because site F486 is directly involved in hACE2-RBD interaction (Shang et al., 2020), explaining at least in part strong selection at this site. Moreover, the branch-site model bsA (positive selection) vs bsA1 (relaxation) of CODEML were compared to find evidence of sites under positive selection in branch of SARS-CoV-2 using the ‘closer’ dataset, but bsA does not show significant differences with bsA1 (p>0.05) indicating selection cannot be confidently implicated, but it was when other datasets were used (including F486). In summary, we do find evidence of sites under positive/episodic selection in branches of close related strains of Wuhan-Hu-1 isolate coronavirus. However, there is not strong evidence of specific sites under positive selection in SARS-CoV-2 using the tools mentioned in this work. This result does not disregard the presence of positive selection sites in SARS-CoV-2, nonetheless, it shows the limitation of the methods to identify with precision specific sites under positive selection in a precise taxon of a phylogenetic tree. We further warn researchers need to be conservative with interpretations of studies utilizing these methodologies, given the equivocal results can be generated by datasets varying in genetic similarity.

To complement our analyses looking for evidence of selection among lineages, we specifically analyzed for patterns of selection across sites in the S-protein genes, we used the sites models available in CODEML and HyPhY. Model M2 of CODEML detected 0.133 % of sites under positive selection (ω>1) and models M1 and M2 detected 85% of sites under purifying selection (ω<1). Model M2 explains the significant data better (p=7e-4) than M1 model, that takes in account only sites with neutral and purifying selection. FUBAR of HyPhy also detected 1070 sites under purifying selection and only 2 sites under positive selection (alignment size 1284). A similar analysis performed by Benvenuto et al. (2020) with FUBAR detects 1065 sites under purifying selection and sites 536 and 644 as under positive selection.

These results suggests by in large strong purifying selection is acting over the vast majority of the S-protein gene, with a comparatively low proportion of sites under positive selection. The evidence of a high amount of purifying selection strengthens similar findings previously reported in RNA viruses (Hughes & Hughes, 2007). These results are congruent with the analysis of Tang et al. (2020) performed with entire coronavirus genomes. However, the sites under positive selection reported differ from our results, and from those of Benvenuto et al. (2020). They found the sites 439, 483 and 439 (counting from the first amino acid of S-protein), our FUBAR analysis found the sites 443 and 445, and FEL found the residue 486 as specific site under positive selection in SARS-COV-2. To resolve these ambiguities in positive selection sites we calculate putative selection sites with CODEML (using Bayes Empirical Bayes from M2 and M8 models) and FUBAR with different datasets reflecting the addition of novel sequences to online repositories (broad, closer, closer without MN996532 and MT084071 and closer without MT084071) and we obtain different results.

It is becoming increasingly clear that predictions of positive selected sites are highly influenced simply by the diversity of the individual sequences included in the datasets. In any case, the majority of predicted sites converge in the region between 439 to 508, a section of the RBD. Additionally, we used TreeSaap to detect important biochemical amino acid properties changes over regions and/or sites along betacoronavirus S-protein. Using a sliding window size of 20 (increasing by 1) we detect that the region between 466 to 500 (using SARS-COV-2 S-protein as a reference) have drastic amino acid changes for alpha-helical tendencies. In addition, the section between 448 to 485 residues registers radical changes in amino acids implicated in the equilibrium constant (ionization of COOH). In the structural analysis we performed, the section between 472 to 486 forms a loop that is not present in certain S-proteins of coronavirus isolated in bats. This loop extends the interaction area between RBD of S-protein and human ACE2, in fact, the lack of these loop decreases the negative energy of interaction (increasing the binding) among these two molecules (see Table 2). These results obtained from independent analysis strongly highlight the importance of 439 to 508 section. Additionally, important hACE2-binding residues in the RBD from SARS-COV-2 obtained from the crystallography and structure determination performed by Shang et al. (2020) are also present in the section we highlight here. We propose that this region is the most probable to contain the sites under positive selection due to predictions by our CODEML and FUBAR models. In that sense, we refer to this section as Region under Positive Selection (RPS). It is important to additionally clarify that even inside the RPS we found at least 20 aa highly conserved between coronaviruses, several of them are predicted as sites under purifying selection. This shows that it is necessary to maintain sites without change around polymorphic sites, probably to conserve the protein structure and at the same time to have the ability to colonize more than one host.

**Table 2.** Estimated free energies of binding of the coronaviruses RBDs to ACE2. Energy values are expressed in kcal/mol.

| RBD | VDWAALS | EEL | EPB | ENPOLAR | EDISPER | ΔG TOTAL |
| --- | --- | --- | --- | --- | --- | --- |
| SARS2 | -89.08 | -180.40 | 180.71 | -64.30 | 129.87 | -23.19 |
| SARS | -88.94 | -171.58 | 175.65 | -64.10 | 133.81 | -15.18 |
| Rs4231 | -80.93 | -139.61 | 141.89 | -56.96 | 119.89 | -15.71 |
| Rm1 | -77.06 | 125.14 | -101.69 | -56.66 | 122.30 | 12.04 |
| SARS2-MUT | -83.88 | -89.47 | 103.76 | -59.49 | 127.43 | -1.66 |

Interestingly, the RPS of the pangolin coronavirus isolate MP789 differs only in one amino acid with the homologous region of SARS-CoV-2, whereas in contrast the bat coronavirus RaTG13 (the overall most similar isolate to SARS-CoV-2 sequenced at the moment) shows 17 differences in the same region. Several explanations could derive from this observation. The hypothesis of recombination inside the pangolin between a native coronavirus strain and a bat coronavirus (like RaTG13) is congruent with our observation. This scenario was proposed and discussed as the origin of SARS-CoV-2 by (Lam et al., 2020; Wong et al., 2020; Xiao et al., 2020), however, other explanations are possible. If the SARS-CoV-2, RaTG13 and pangolin coronavirus MP789 isolate are closely related as shown in the tree of the Figure 1, we are observing the ancestral sequence of RPS in human and pangolin coronaviruses, and a mutated version in bat virus. Elucidating the origin of SARS-CoV-2 is beyond the scope of this work, nevertheless sequencing of new coronavirus isolates in the near future could resolve this question.

With a list of broader observations related to the role of selection across viral genomes we aimed to specifically understand how these regions could affect virus/host interactions. To understand more in deep the importance of RPS in the evolution of SARS-COV-2, we quantified the relative importance of this region in the interaction between RBD and hACE2. In that sense, MD simulations were run for five complexes (listed in methods). In all cases the systems were stable with Root Mean Square Deviations (RMSD) of their backbones between 1.11 Å and 3.30 Å relative to the initial complexes structures during the last 5 ns of the production run. We first investigated the network of contacts between the ligands (coronaviruses RDB) with the receptor (hACE2). Overall, all complexes present a large number of contacts between the ligands and the receptor in at least 50% of the MD snapshots selected for MM-PBSA calculations. Common interactions with T27, F28, K31, H34, Y41, K353, G354, D355 and R357 of the receptor are observed in all systems. The full networks of interactions between the coronaviruses and the hACE2 receptor are provided as Supporting Information.

Next we estimated the free energies of binding of the coronaviruses’ RBDs to hACE2 and the results of these evaluations are summarized in Table 2. These calculations show that the SARS2, SARS and Rs4231 viruses are predicted to favorably bind to the human hACE2 receptor, while the Rm1 and SARS2-MUT variants present unfavorable free energies of binding. The fact that the bat’s coronavirus Rs4231, in addition to SARS and SARS2, presents favorable interaction with hACE2 is in accordance with the previous observation that it is able to infect human cells expressing this protein (Hu et al., 2017). To get more insights into the contribution of the receptor and the RBDs to the binding process, we performed energy decomposition experiments.

The contribution of each residue in the studied coronaviruses that interact with the hACE2 receptor are shown in Table 3. Rows are presented in such a way that each of them contains the residues occupying the same position in the viruses RBDs structures as in the SAR2 RBD structure. From here on, residues numeration will take that of SARS2 as reference. In general, most RBDs residues show negative values of contribution to the free energies of binding to the human receptor. All studied RBDs, except that of the Rm1 coronavirus, have amino acids with large favorable contributions to the free energies of binding that directly interact with hACE2: K417 of SARS2 and SARS2-MUT, R426 in SARS and R480 in Rs4231. On the other hand, the G476D mutation (D463 present in bat coronavirus strains) have a negative contribution to the binding of the RDB to hACE2. This site was predicted to be under purifying selection by FUBAR analyses, and is located within the RPS.

**Table 3.** Contribution of the residues in the RDB of the coronaviruses interacting with the hACE2 receptor. Energy values are presented as kcal/mol.

| SARS2 |  | SARS |  | Rs4231 |  | Rm1 |  | SARS2-MUT |  |
| --- | --- | --- | --- | --- | --- | --- | --- | --- | --- |
| K417 | -93.13 | V404 | -2.16 | V405 | -1.70 | V408 | -1.66 | K417 | -98.98 |
| N439 | -0.61 | R426 | -127.22 | N427 | -0.82 | A430 | 0.29 | N439 | -0.40 |
| Y449 | -8.28 | Y436 | -4.85 | Y437 | -1.53 | Del. | 0.00 | Y449 | -1.19 |
| Y453 | -1.33 | Y440 | -1.77 | Y441 | 1.04 | Y439 | -0.69 | Y453 | -1.60 |
| L455 | -1.38 | Y442 | -4.11 | W443 | -1.93 | S441 | -1.75 | L455 | -1.43 |
| F456 | -3.50 | L443 | -1.62 | V444 | -0.74 | Y442 | -3.53 | F456 | -2.70 |
| A475 | -5.05 | P462 | -4.05 | P463 | -7.75 | Del. | 0.00 | A475 | -3.92 |
| G476 | -2.82 | D463 | 58.07 | G464 | -3.45 | Del. | 0.00 | G476 | -1.66 |
| F486 | -3.58 | L472 | -3.37 | P473 | -2.04 | Del. | 0.00 | F486 | -5.03 |
| N487 | -6.31 | N473 | -4.08 | N474 | -1.89 | N459 | 0.05 | N487 | -4.55 |
| Y489 | -3.52 | Y475 | -3.32 | Y476 | -2.75 | V461 | -3.52 | Y489 | -2.59 |
| F490 | 0.30 | W476 | 0.87 | N477 | 0.41 | Y462 | -4.09 | F490 | 0.55 |
| L492 | -1.26 | L478 | 0.77 | L479 | 0.25 | L464 | -3.24 | L492 | -0.78 |
| Q493 | -14.68 | N479 | -5.86 | R480 | -147.75 | S465 | -3.84 | Q493 | -18.36 |
| Y495 | -2.17 | Y481 | -0.73 | Y482 | -2.35 | Y467 | -4.28 | Y495 | -1.70 |
| G496 | -2.21 | G482 | -0.72 | G483 | -1.50 | D468 | 105.79 | D496 | 103.96 |
| Q498 | -4.67 | Y484 | -3.98 | F485 | -3.27 | Y470 | -10.31 | Q498 | -3.45 |
| T500 | -4.90 | T486 | -6.96 | T487 | -6.61 | S472 | -8.38 | T500 | -4.80 |
| N501 | -12.17 | T487 | -6.79 | A488 | -4.35 | I473 | -5.41 | N501 | -1.69 |
| G502 | -4.21 | G488 | -3.48 | G489 | -3.25 | P474 | -2.97 | G502 | -3.89 |
| Y505 | -5.99 | Y491 | -7.16 | H492 | -7.11 | Y477 | -10.14 | Y505 | -7.77 |

Strikingly, the G496D mutation (SARS2 numeration) has a large negative influence in the free energy of binding in the two complexes that contain it. It is also worth noting that the three aspartic acid substitutions present in all systems negatively contribute to the systems stability. Taking into account that the only difference between SARS2 and SARS2-MUT is the G496D mutation, we postulate that this RBD position is critical for the human receptor recognition by coronaviruses. To the best of our knowledge, no coronavirus having aspartic acid at this position is able to infect human cells. This result supports the prediction from FUBAR analyses indicating that the site G496D is under purifying selection. Combined, our results strongly suggest that the mutation of the D496 residue present in the coronaviruses from bats is critical for their RBDs to recognize the human hACE2 receptor. Additionally, it shows the importance of sites under purifying selection in RPS for the RBD evolution.

To better interpret the influence of the key interactions between the coronaviruses RBDs and their hACE2 receptor, their interactions were analyzed. To select the representative structure of each system the MD snapshots employed for MM-PBSA calculations were clustered. Then, the representative structure of a system was selected as the centroid of the most populated cluster. The predicted RBD-hACE2 complexes for SARS2, SARS and SARS2-MUT are depicted in Figure 2.

**Figure 2.**
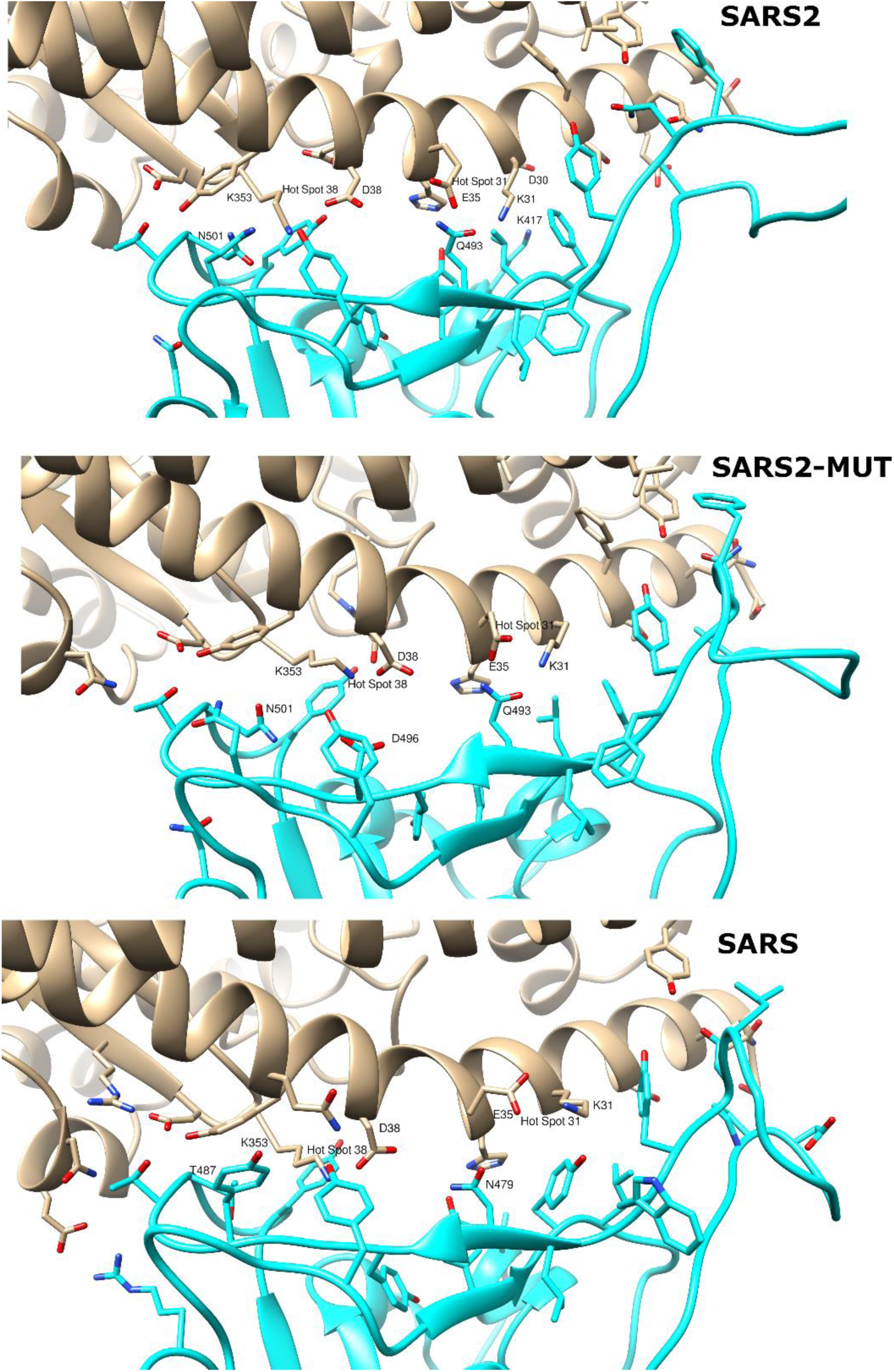
Predicted interaction of SARS2 (top), SARS2-MUT (middle) and SARS (bottom) with the human ACE2 receptor. hACE2 in depicted in gray and the RBD of coronaviruses in cyan. Oxygen atoms are colored red and nitrogen blue. The numbering of the residues corresponds to that of the sequence of each spike glycoprotein.

Many studies have focused on coronaviruses mutations that favor adaptations for human hosts infections. For example, it has been shown that specific substitutions at positions 455, 486, 483, 494 and 501 (442, 472, 479, 480 and 487 in SARS) of the RBD of SARS favors the interaction between the RBD of SARS and hACE2 (Cui et al., 2019). Likewise, homology modeling studies found favorable interactions between the residues occupying these positions in the SARS2 RBD and the human receptor (Wan et al., 2020). The cornerstone of these favorable interactions is the complementarity of the RBDs with hot spots 31 and 353. These are salt bridges between K31 and E35 and between D38 and K353 of ACE2 which are buried in a hydrophobic environment (see Figure. 2). In the cases of SARS2 and SARS, Q493 (N479 in SARS) and N501 (T487 in SARS) add support to the hot spots according to these previous studies. These observations should also hold for the Rs4231 strain, however the N501A change in the later compared to SARS2 (A488 in Rs4231) add little support to hot spot 353. In this case, to continue permitting human infection, the large favorable contribution of R480 in Rs4231 to the free energy of binding could compensate the weak support provided by A488 to hot spot 353.

Interestingly, K353 is the residue forming the largest network of contacts with the analyzed RBDs among those belonging to both hot spots. Our simulations also show that in SARS2 and SARS the RBD amino acids with the largest contribution to the free energy of binding, K417 and R426 (see Table 3) respectively, do not interact with any hot spot residue. Instead, they interact with D30 of hACE2 in the SARS2 complex and with E329 of the human receptor in the SARS complex. This could indicate that interactions additional to those previously identified with the hACE2 hotspots could be critical for the stabilization of the RDB-human receptor complexes. Finally, we analyzed the possible reasons for the predicted negative impact that the G496D mutation has on the predicted free energies of binding of the RBD to hACE2. As depicted in Figure 2, G496 directly interacts with K353 in hot spot 353 and its mutation interferes with the D38-K353 salt bridge. Specifically, D496 of the RDB point to D38 of hACE yields a high electric repulsion between these amino acids. Consequently, this portion of the RBD is pushed to a position further from hACE2 than that observed in the wild type receptor, resulting in the reduction of its network of contacts with K353. As a result, the binding of the RBD to hACE2 is considerably inhibited and unlikely to occur.

## Conclusions

A priority in ongoing research is to better understand coronavirus evolution, with specific interests in understanding the role of selection pressures in viral evolution, and clarifying how viral strains can infect novel hosts. Our experiments suggest that there are sites under positive selection in the S-protein gene of SARS-CoV-2 and other betacoronaviruses, particularly in a region that we called RPS (Region under Positive Selection) inside of the RBD. However, we have identified that by in large, sites in this region (and overall, in the S-protein gene) are under purifying selection. Particularly, for the site D496G, the presence of aspartic acid seems indispensable for the interaction with the hACE2. Additionally, we performed MD simulations and free energies of binding predictions for five different complexes of coronaviruses that do and do not infect human cells. Our results suggest that as long as no disrupting interference occur with both salt bridges at hot spots 31 and 353 coronaviruses are able to bind with hACE2. Modeling results suggest that interference with the hot spot 353 could be and effective strategy for inhibiting the recognition of the RBD of the SARS-COV-2 spike protein by its human host receptor ACE2 and hence prevent infections. Although additional simulations and experiments are required, all evidence suggests that the mutation of D496 in the bat variants of the coronaviruses permit infection of human cells. Giving the large contribution of SARS2 K417 to the free energy of binding of the RBD to hACE2 we propose that blocking its interaction with the receptor D30 could be a promising strategy for future drug discovery efforts.

