## Supporting information for "SARS-CoV-2, an evolutionary perspective of interaction with human ACE2 reveals undiscovered amino acids necessary for complex stability"


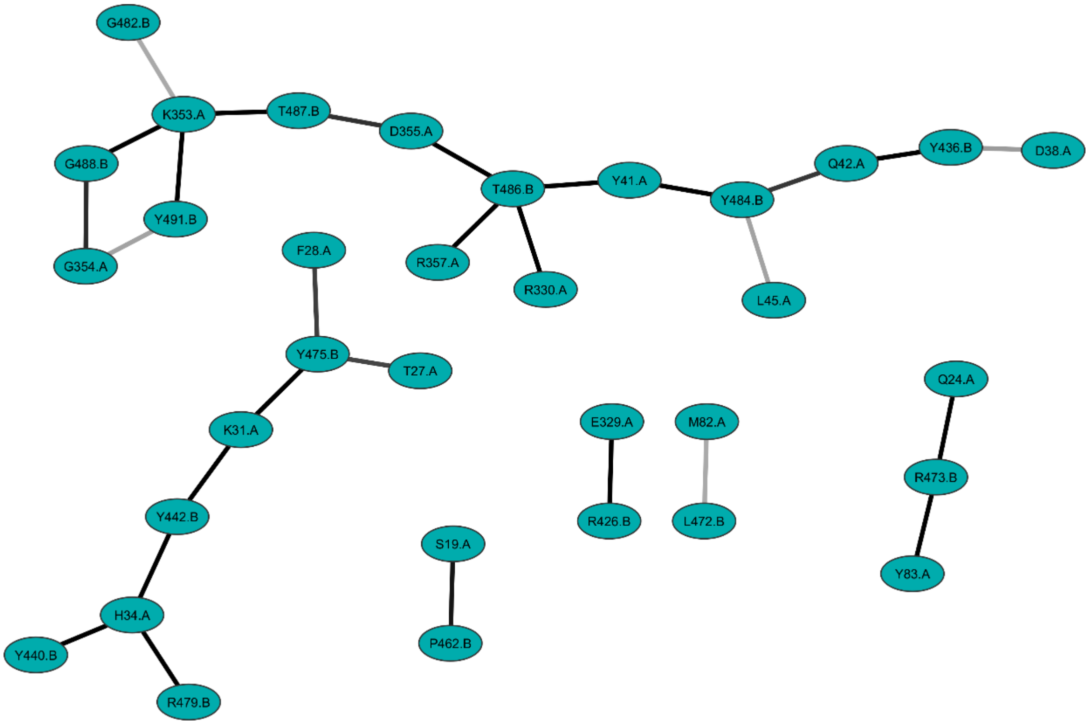


*Figure S1. Network of contacts between the hACE2 receptor and the RBD of the SARS coronavirus along the MD simulation. Darker lines indicate more frequent interactions. Residues are labelled with "A" and "B" to indicate whether they belong to the human receptor or the virus RBD, respectively.*

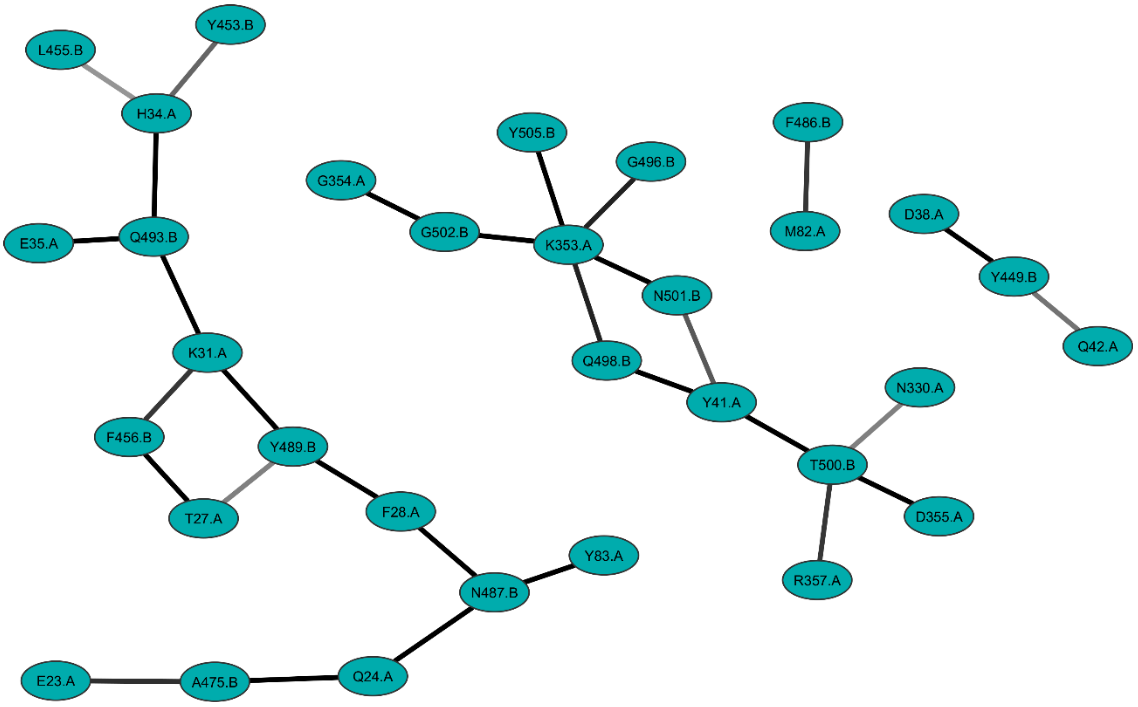


*Figure S2. Network of contacts between the hACE2 receptor and the RBD of the SARS2coronavirus along the MD simulation. Darker lines indicate more frequent interactions. Residues are labelled with "A" and "B" to indicate whether they belong to the human receptor or the virus RBD, respectively.*

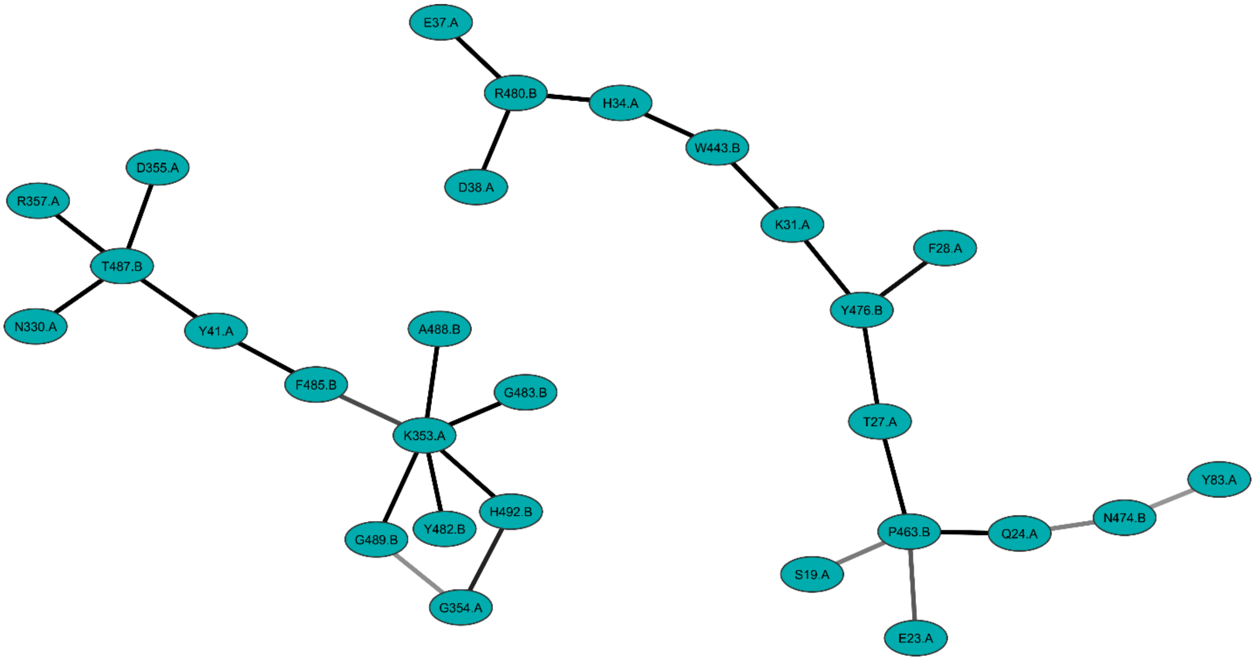


*Figure S3. Network of contacts between the hACE2 receptor and the RBD of the Rs4231 coronavirus along the MD simulation. Darker lines indicate more frequent interactions. Residues are labelled with "A" and "B" to indicate whether they belong to the human receptor or the virus RBD, respectively.*

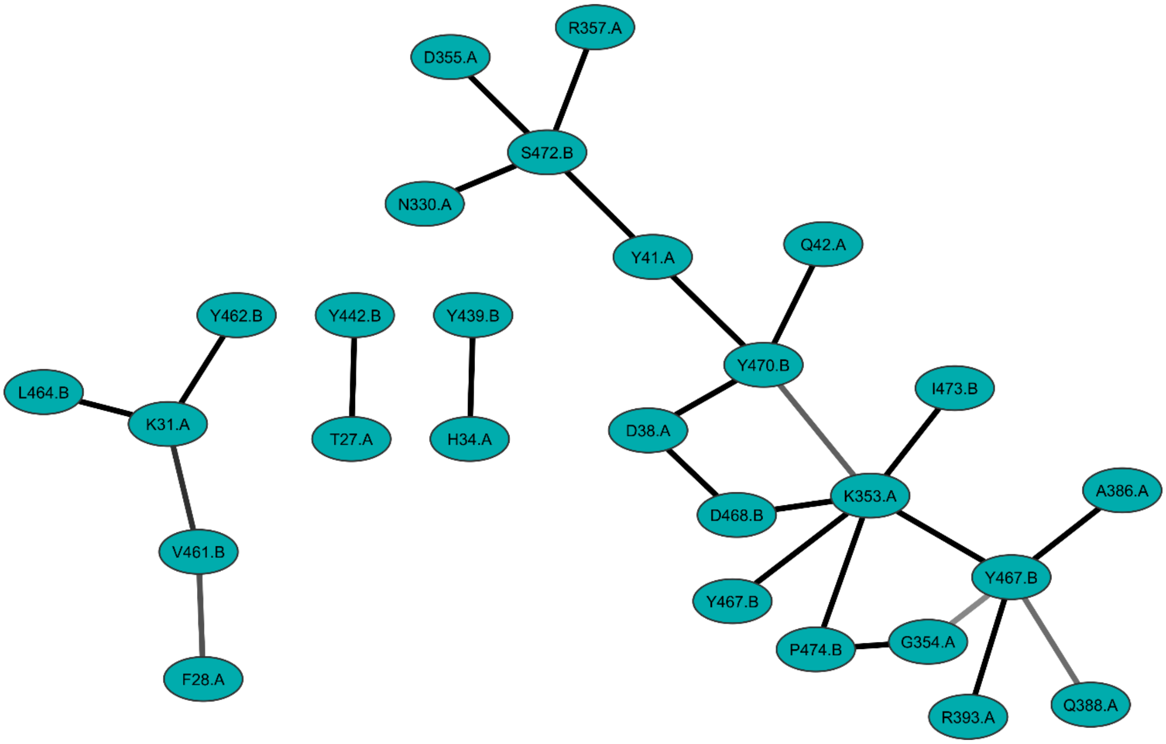


*Figure S4. Network of contacts between the hACE2 receptor and the RBD of the Rm1 coronavirus along the MD simulation. Darker lines indicate more frequent interactions. Residues are labelled with "A" and "B" to indicate whether they belong to the human receptor or the virus RBD, respectively.*

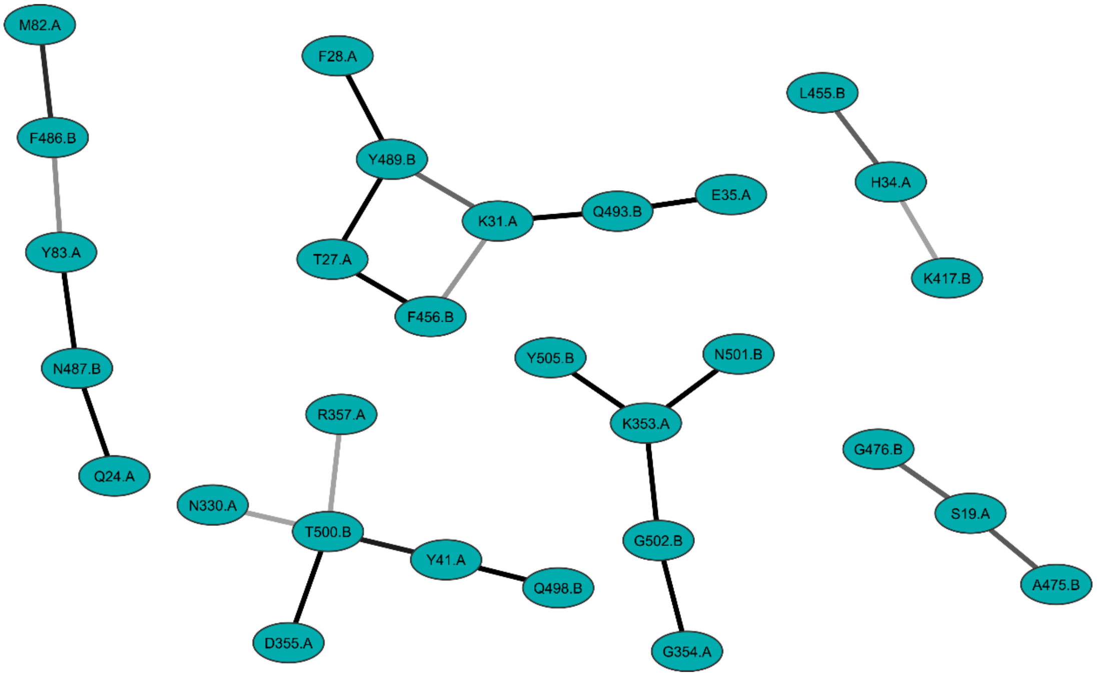


*Figure S5. Network of contacts between the hACE2 receptor and the RBD of the SARS2-MUT coronavirus along the MD simulation. Darker lines indicate more frequent interactions. Residues are labelled with "A" and "B" to indicate whether they belong to the human receptor or the virus RBD, respectively.*
